# Mouse microglia express unique miRNA-mRNA networks to facilitate age-specific functions in the developing central nervous system

**DOI:** 10.1101/2022.07.12.499835

**Authors:** Alexander D. Walsh, Sarrabeth Stone, Andrea Aprico, Trevor J. Kilpatrick, Brendan A. Ansell, Michele D. Binder

## Abstract

Microglia regulate multiple processes in the central nervous system, exhibiting a significant level of cellular plasticity which is facilitated by an equally dynamic transcriptional environment. While many gene networks that regulate microglial functions have been characterised, the influence of epigenetic regulators such as small non-coding microRNAs (miRNAs) is less well defined. We have sequenced the miRNAome and mRNAome of mouse microglia during brain development and adult homeostasis, identifying unique profiles of known and novel miRNAs. Microglia express both a consistently enriched miRNA signature as well as temporally distinctive subsets of miRNAs. We generated robust miRNA-mRNA networks related to fundamental developmental processes, in addition to networks associated with immune function and dysregulated disease states. There was no apparent influence of sex on miRNA expression. This study reveals a unique developmental trajectory of miRNA expression in microglia during critical stages of CNS development, establishing miRNAs as important modulators of microglial phenotype.

## Introduction

Microglia are dynamic regulators of the central nervous system (CNS) where they support early development, adult homeostasis and immune function. Following embryonic colonization of the brain microglia regulate neurogenesis, synaptogenesis and myelination via both the clearance (efferocytosis) of immature cells and synapses, and secretion of trophic factors^1–4^. In the adult brain microglia continue to regulate existing neural networks, as well as support neurogenic niches and oligodendrocyte precursor cell pools^5,6^. Adult microglia adopt a ramified phenotype and extend cellular processes to survey the local environment and maintain tissue homeostasis^7,8^. Upon interaction with pathological stimuli, microglia can rapidly shift their phenotype to initiate apoptotic clearance and cytokine signaling to suppress inflammation and promote a neuroprotective environment^9^.

Given the broad spectrum of phenotypes and activities of microglia in the healthy CNS, appropriate microglial function is heavily dependent upon precise regulation of gene expression. Consequently, genetic dysregulation of microglial biology is implicated in numerous neurological disorders. Chronically activated ‘neurotoxic’ microglia have been identified as a hallmark of neurodegeneration and autoimmunity, with key genes and signaling pathways explicitly linked to the pathology of multiple sclerosis (MS), amyotrophic lateral sclerosis (ALS), Alzheimer’s disease (AD) and Huntington’s disease (HD)^10^. Perturbed microglial activity is also implicated in neurodevelopmental disorders including autism spectrum disorder (ASD), schizophrenia and epilepsy^11–14^.

Multiple sequencing studies of microglial populations have captured genetic and epigenetic networks that regulate cell identity and define microglial phenotypes in the healthy and diseased brain^15–19^. In addition, comprehensive profiling has identified distinct programs of gene expression that tightly regulate developmental phenotypes and age-specific microglial functions^20,21^. Pre-natal and early post-natal microglia are highly reactive and adopt an ameboid morphology similar to that of adult activated microglia during inflammation and disease. However, there is no strong gene expression overlap between these cell subsets, indicating that developmental microglia are a distinct cell population that uniquely contribute to CNS development. These studies highlight the multiplicity of transcriptional programmes which can be activated by microglia in response to normal development or pathological challenge. An important question then arises of how are these transcriptional programmes controlled? One strong potential candidate is the class of small RNAs known as microRNAs (miRNAs).

miRNAs are small (18-22bp) non-coding RNAs that act as negative post-transcriptional regulators of gene expression^22^. Mature miRNAs are integrated into a catalytic complex that binds to target mRNAs and trigger degradation or stalled translation of the target mRNA transcript^23^. Ubiquitous in the mammalian genome, miRNAs regulate fundamental cellular processes including cell differentiation, proliferation and immune homeostasis^24,25^. Tissue specific sequencing of miRNAs has revealed that ubiquitously expressed miRNAs are less common than previously predicted and emphasise the importance of cell specific studies to capture accurate miRNA profiles^26^. Normal specification of microglia is known to be reliant upon miRNAs, as conditional knockout of Dicer in microglia, which ablates the miRNA biogenesis pathway, results in perturbation of microglia and induces hyper-responsiveness to inflammatory stimuli^27^. In addition, specific miRNAs have a profound influence on microglial biology. For example, miR-155 and miR-124 have been identified as ‘master regulators’ of activated and quiescent microglia respectively^28–30^. However, much of the evidence for miRNA mediated regulation of microglia stems from studies of adult mice, and thus the role of miRNAs in regulating developmental microglial functions are not as well understood. Further, studies of the miRNAome are often limited in their ability to predict transcriptional effects, as they rely on *in silico* predictions of miRNA-mRNA interactions rather than integrated network analyses.

In this study we have characterised the miRNAome of microglia in male and female mice at three developmental timepoints; post-natal days 6, 15, and 8 weeks. We identified a unique microglial miRNA profile compared with other CNS cell types. Differential expression analysis identified age-specific subsets of miRNA expression, suggesting a gene regulatory programme that influences microglial biology during development. Unexpectedly, we did not observe any sex-specific differential miRNA expression, and found only a modest difference in mRNA expression, indicating that microglia are not necessarily innately different between sexes during development. Integrated miRNA-mRNA expression analyses identified 266 mRNAs that were negatively correlated with enriched microglial miRNAs, and that were predicted to target key neurodevelopmental processes including neurogenesis, axonal guidance and myelination. These data provide a unique profile of the microglial transcriptome and its epigenetic regulation, providing critical insight into the role of microglial miRNAs in regulating microglial developmental programmes.

## Results

### Isolation of highly pure microglial populations via CD45+ve immunopanning

To explore the role of microglial enriched miRNAs and their influence on gene regulatory networks during CNS development, we sequenced and analysed the miRNA and mRNA transcriptomes of CD45^+ve^ microglia isolated from three age groups of C57Bl/6 mice (P6, P15, 8 weeks) as well as the miRNA expression of the CD45^-ve^ bulk CNS cells (primarily comprised of neurons, oligodendrocytes and astrocytes) (Fig. 1). Expression analyses demonstrated that microglial specific markers such as Tmem119 and Cx3cr1 were highly expressed by purified CD45^+ve^ cells (Fig. S1).

**Figure 1:**
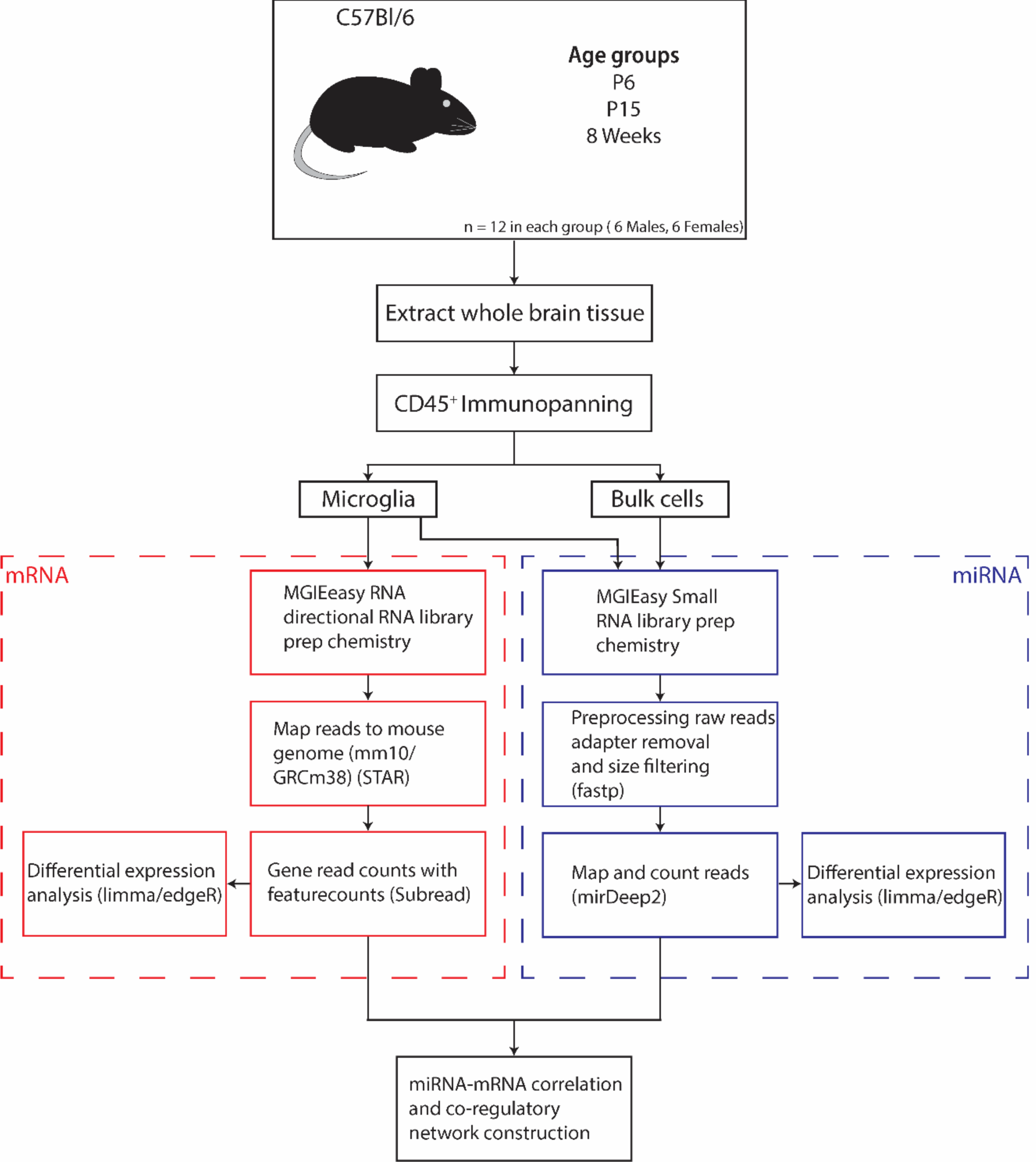
miRNA/mRNA sequencing and analysis workflow. Whole brains of 3 groups of C57Bl/6 mice (n=12) were processed and CD45^+ve^ microglia were isolated via immunopanning. Microglia and bulk cell populations were sequenced for miRNA expression and analysed for differential expression using the edgeR/limma pipeline. In addition, mRNA expression was sequenced in microglial populations and read counts were correlated with miRNA data to construct miRNA-mRNA co-regulatory networks.

Multidimensional scaling (MDS) analysis of miRNA sequencing libraries identified strong separation of sample populations by both cell type (microglia vs bulk) as well as age (Fig. 2A). Overall, 1075 unique miRNAs were expressed across all mice, with a minimum count per million (CPM) of 1 in at least 5 samples. Sequencing of microglial mRNA transcriptomes identified 15,660 robustly expressed transcripts (minimum CPM of 0.5 in at least 5 samples) and MDS analysis revealed a similarly strong separation of samples by age (Fig. 2B). Amongst the most highly expressed microglial miRNAs and mRNAs were genes previously associated with microglial identity and homeostasis, including *Cx3cr1, Csf1r, Hexb, Sparc*, miR-124, let-7 and miR-29b, further validating the microglial enrichment of the CD45^+ve^ sequencing population (Table 1).

**Table 1.**
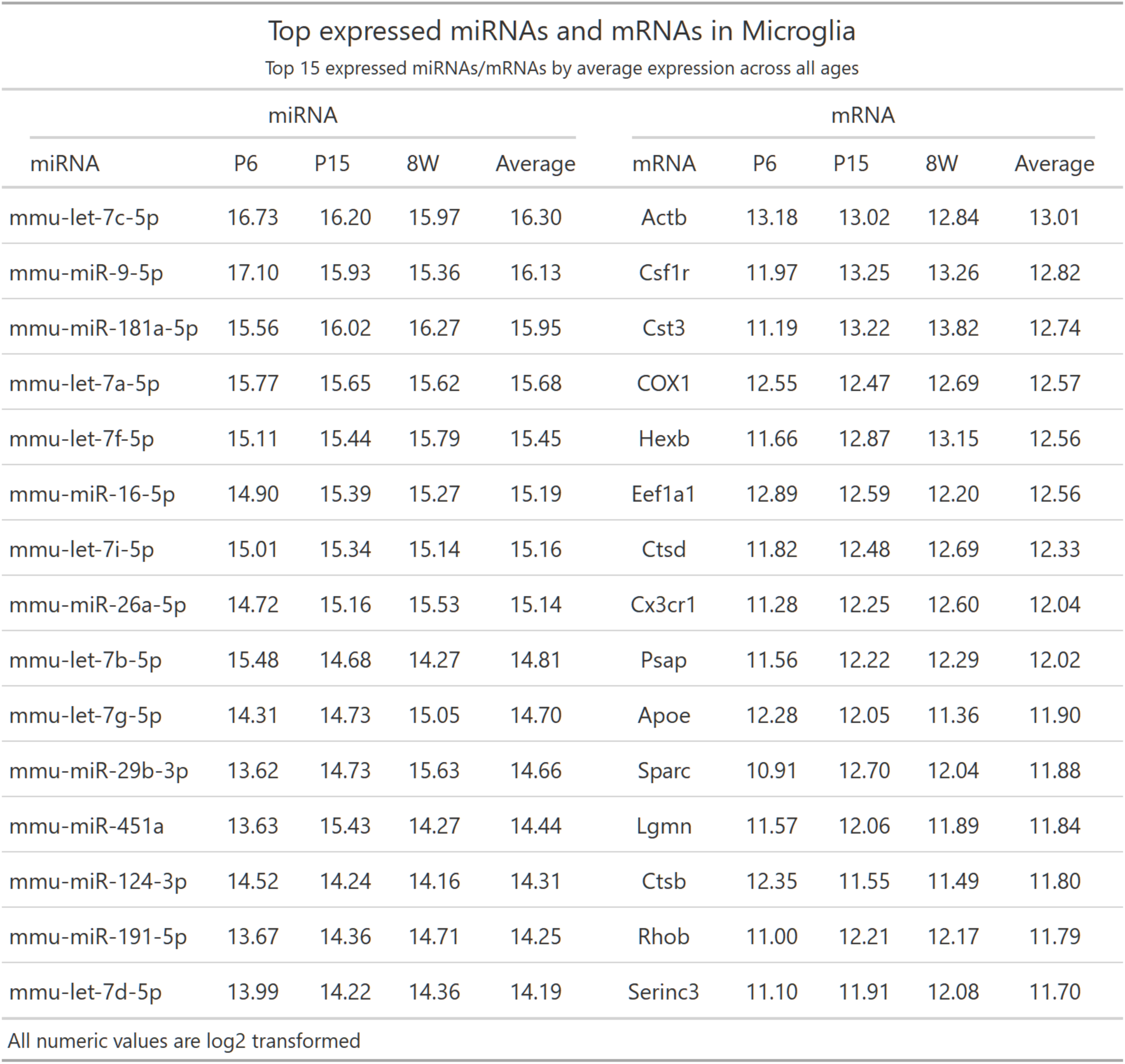

| Top expressed miRNAs and mRNAs in Microglia |  |  |  |  |  |  |  |  |  |
| --- | --- | --- | --- | --- | --- | --- | --- | --- | --- |
| Top 15 expressed miRNAs/mRNAs by average expression across all ages |  |  |  |  |  |  |  |  |  |
| miRNA |  |  |  |  | mRNA |  |  |  |  |
| miRNA | P6 | P15 | 8W | Average | mRNA | P6 | P15 | 8W | Average |
| mmu-let-7c-5p | 16.73 | 16.20 | 15.97 | 16.30 | Actb | 13.18 | 13.02 | 12.84 | 13.01 |
| mmu-miR-9-5p | 17.10 | 15.93 | 15.36 | 16.13 | Csf1r | 11.97 | 13.25 | 13.26 | 12.82 |
| mmu-miR-181a-5p | 15.56 | 16.02 | 16.27 | 15.95 | Cst3 | 11.19 | 13.22 | 13.82 | 12.74 |
| mmu-let-7a-5p | 15.77 | 15.65 | 15.62 | 15.68 | COX1 | 12.55 | 12.47 | 12.69 | 12.57 |
| mmu-let-7f-5p | 15.11 | 15.44 | 15.79 | 15.45 | Hexb | 11.66 | 12.87 | 13.15 | 12.56 |
| mmu-miR-16-5p | 14.90 | 15.39 | 15.27 | 15.19 | Eef1a1 | 12.89 | 12.59 | 12.20 | 12.56 |
| mmu-let-7i-5p | 15.01 | 15.34 | 15.14 | 15.16 | Ctsd | 11.82 | 12.48 | 12.69 | 12.33 |
| mmu-miR-26a-5p | 14.72 | 15.16 | 15.53 | 15.14 | Cx3cr1 | 11.28 | 12.25 | 12.60 | 12.04 |
| mmu-let-7b-5p | 15.48 | 14.68 | 14.27 | 14.81 | Psap | 11.56 | 12.22 | 12.29 | 12.02 |
| mmu-let-7g-5p | 14.31 | 14.73 | 15.05 | 14.70 | Apoe | 12.28 | 12.05 | 11.36 | 11.90 |
| mmu-miR-29b-3p | 13.62 | 14.73 | 15.63 | 14.66 | Sparc | 10.91 | 12.70 | 12.04 | 11.88 |
| mmu-miR-451a | 13.63 | 15.43 | 14.27 | 14.44 | Lgmn | 11.57 | 12.06 | 11.89 | 11.84 |
| mmu-miR-124-3p | 14.52 | 14.24 | 14.16 | 14.31 | Ctsb | 12.35 | 11.55 | 11.49 | 11.80 |
| mmu-miR-191-5p | 13.67 | 14.36 | 14.71 | 14.25 | Rhob | 11.00 | 12.21 | 12.17 | 11.79 |
| mmu-let-7d-5p | 13.99 | 14.22 | 14.36 | 14.19 | Serinc3 | 11.10 | 11.91 | 12.08 | 11.70 |
| All numeric values are log2 transformed |  |  |  |  |  |  |  |  |  |

**Figure 2:**
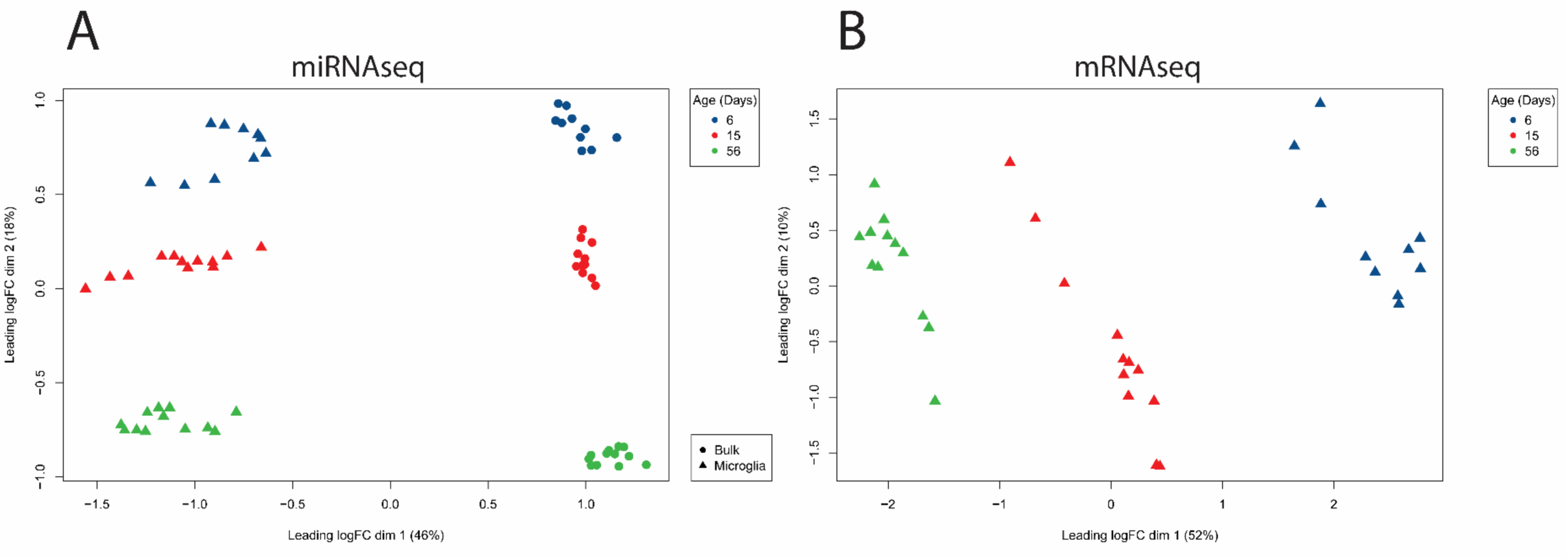
MDS analysis of isolated samples identifies strong clustering by sample age and cell type status. MDS clustering of **A**. miRNA sequencing data (microglia and bulk samples) and **B**. mRNA sequencing data (microglia only) was performed for 34 mice. miRNA samples clustered by both age (in days) and cell type. Similarly, mRNA sequenced microglia samples were strongly separated by developmental age.

### Developmental microglia express a common miRNA signature and subsets of temporally dynamic miRNAs

A primary aim of this study was to characterise miRNAs enriched in microglia relative to other CNS cell types. To do this, we first undertook a general miRNA enrichment analysis (pooling and comparing all microglial samples to all bulk samples, controlling for age, sex and mouse ID) which identified 244 miRNAs as upregulated in microglia (Fig. 3A). The most highly enriched miRNAs included many that were previously defined as important in regulating microglial identity, including miR-142a-5p, miR-223-3p and miR-146a-3p (Fig. 3A). Also identified as highly enriched were miR-1895, miR-511-3p and miR-6983-5p, which have less well-defined functions in microglia (Fig. 3A). Next, we performed the enrichment analysis within each age group to generate age-specific miRNAomes (Table S1). Three hundred and twenty-nine miRNAs were defined as enriched in at least one age group, with 91 enriched at all ages, indicative of a common signature of microglial miRNA expression (Fig. 3B). The top 10 enriched miRNAs in each age group are presented in Table 2. Conversely, 129 miRNAs were enriched in a single age group, with 66 of these specific to adult cells (Fig. 3B). To validate expression of top enriched candidates, we performed qPCR of selected miRNAs, confirming strong upregulation of four miRNAs in 8 week mice; miR-223, miR-146a-3p, miR-455-3p, miR-513p (Fig. 3C). Extended qPCR analysis also validated specific patterns of observed temporal miRNA expression, including examples of candidates enriched across all ages (miR-21a-3p), enriched specifically at P6 and P15 (miR-29a-3p, miR-29b-3p, miR-3473b) and exhibiting increased expression (and enrichment) throughout development (miR-10a-5p) (Fig. 3D). Overall, this enrichment analysis identified commonly enriched as well as age-specific miRNAs that contribute to the developmental miRNAome of microglia.

**Table 2:**

| miRNA enrichment in microglia |  |  |  |  |  |  |  |  |
| --- | --- | --- | --- | --- | --- | --- | --- | --- |
| Top 10 upregulated miRNAs vs. other CNS cell types in each age group |  |  |  |  |  |  |  |  |
| P6 |  |  | P15 |  |  | 8 Weeks |  |  |
| miRNA | Enrichment<sup>1</sup> | Avg Expression<sup>2</sup> | miRNA | Enrichment<sup>1</sup> | Avg Expression<sup>2</sup> | miRNA | Enrichment<sup>1</sup> | Avg Expression<sup>2</sup> |
| mmu-miR-511-3p | 9.77 | 7.51 | mmu-miR-511-3p | 8.04 | 7.34 | mmu-miR-6983-3p | 7.52 | 5.80 |
| mmu-miR-6983-3p | 8.14 | 5.58 | mmu-miR-6983-3p | 7.46 | 6.26 | mmu-miR-223-3p | 7.10 | 11.02 |
| mmu-miR-223-3p | 8.05 | 11.31 | mmu-miR-223-3p | 7.38 | 11.13 | mmu-miR-146a-5p | 6.82 | 14.87 |
| mmu-miR-223-5p | 7.99 | 4.24 | mmu-miR-223-5p | 7.02 | 4.12 | mmu-miR-223-5p | 6.62 | 3.75 |
| mmu-miR-511-5p | 7.87 | 4.38 | mmu-miR-511-5p | 6.99 | 4.20 | mmu-miR-511-3p | 6.42 | 5.63 |
| mmu-miR-1895 | 7.29 | 5.83 | mmu-miR-6983-5p | 6.37 | 3.32 | mmu-miR-142a-5p | 6.30 | 10.97 |
| mmu-miR-142a-3p | 6.86 | 13.75 | mmu-miR-142a-3p | 6.26 | 14.22 | mmu-miR-142a-3p | 6.19 | 13.61 |
| mmu-miR-142a-5p | 6.37 | 10.36 | mmu-miR-142a-5p | 5.96 | 11.06 | mmu-miR-6983-5p | 6.03 | 2.91 |
| mmu-miR-3079-3p | 6.15 | 2.91 | mmu-miR-146a-3p | 5.75 | 4.40 | mmu-miR-146a-3p | 5.97 | 4.97 |
| mmu-miR-6983-5p | 6.15 | 2.47 | mmu-miR-146a-5p | 5.69 | 14.05 | mmu-miR-7219-3p | 5.64 | 4.37 |

<sup>1</sup> Log2 fold change comparing microglia to bulk CNS cell expression
<sup>2</sup> Average expression of miRNA in all microglial samples at designated age
All numeric values are log2 transformed

**Figure 3.**
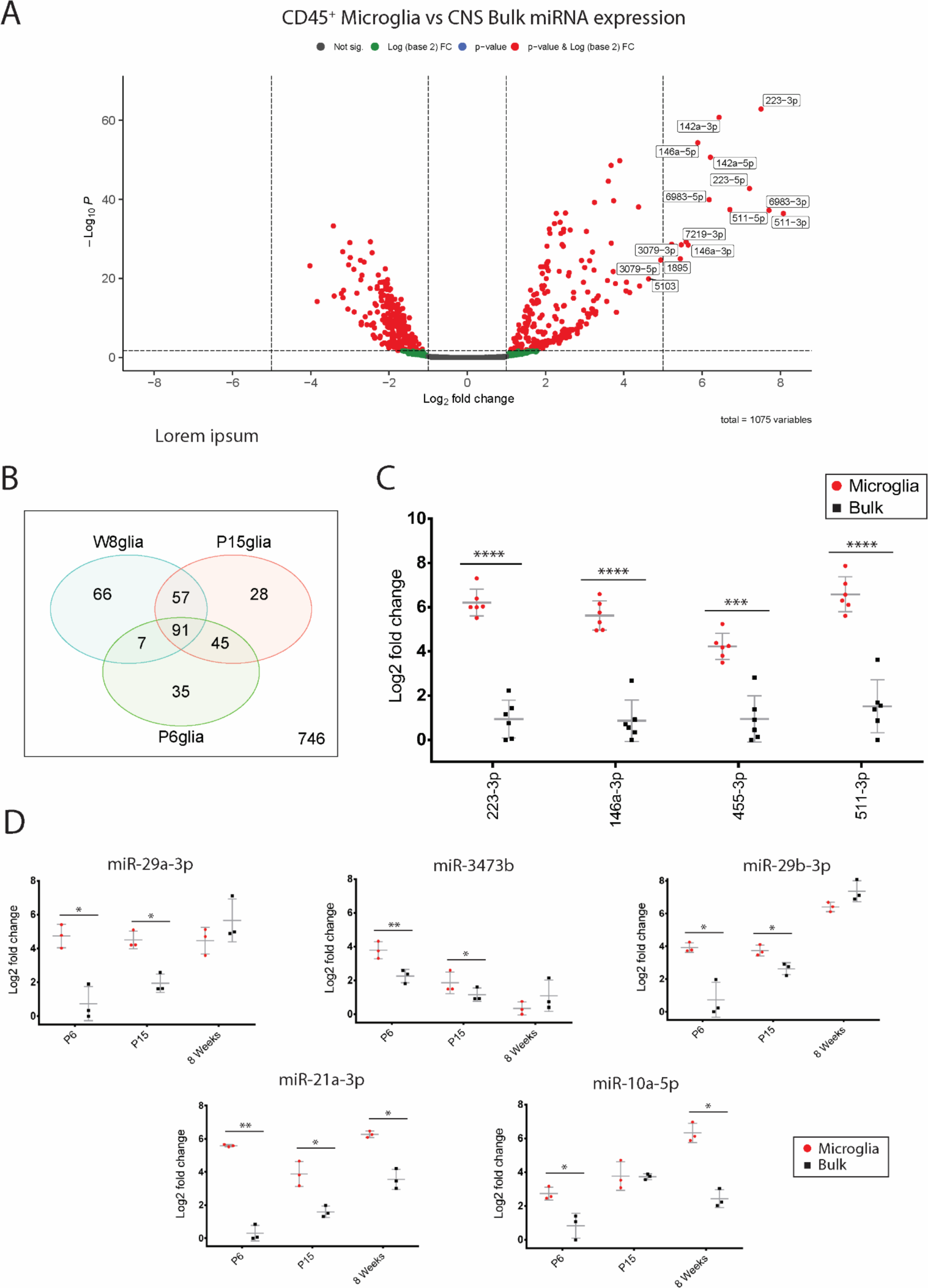
Microglia express unique developmental profiles of miRNA expression compared to surrounding bulk tissue. **A**. Volcano plot of differential miRNA expression between pooled microglia samples and bulk samples correcting for age and sex. 224 of 1075 miRNAs were observed to be upregulated in microglia (logFC >1, FDR < 0.05). Top 15 upregulated miRNAs (by log2 fold change) in microglia are labelled. **B**. Venn diagram comparing subsets of enriched miRNAs (microglia vs bulk) in each age group (logFC >1, p<0.05). 329 miRNAs were observed as upregulated in at least one age group, with 91 consistently upregulated at every age. **C**. qPCR validation of selected miRNA candidates in 8 week old mice (n=6 in each group). Data were analysed by paired t tests comparing microglia and bulk group expression for each tested miRNA.. **D**. qPCR validation of selected miRNA candidates across 3 age groups, P6, P15 and 8 weeks (n = 3 in each group). Data were analysed by paired t test comparing microglia and bulk expression within each age group. * p < 0.05, ** p < 0.01, *** p< 0.001, **** p < 0.0001. Lines on qPCR plots represent mean and standard deviation.

### mRNA-miRNA regulatory network analysis identifies age-specific microglial biology that contributes to CNS development

To characterise the gene regulatory networks potentially regulated by age-specific miRNAomes in microglia, we correlated the expression of microglial enriched miRNAs and all 15,660 expressed mRNA transcripts from isolated microglia populations (Figs. 4,5,6). Given that miRNAs are canonically negatively correlated with their mRNA targets, we selected putative miRNA-mRNA pairs that were strongly negatively correlated (r < -0.894, FDR < 0.05). miRNA-mRNA pairs were further filtered for experimental evidence of interaction as reported in existing literature and relevant databases (collated by multiMiR) and constructed into ‘high confidence’ miRNA-mRNA regulatory networks for each age group. A consistent observation in all these networks was a high degree of interconnectedness, with many genes being targeted by more than one miRNA (Figs. 4A, 5A, 6A). Fourteen miRNAs were present in all age-group networks, creating significant overlap in represented pathways. Conversely, three miRNAs were present in a single network; miR-210-3p at P6, and miR-10a-5p and miR-103-3p at 8 weeks (Figs. 4A, 6A). KEGG pathway enrichment analysis of target mRNAs in each network identified strong representation of biological processes related to cell cycle, neurogenesis, axon guidance and pathways including p53, Hippo, PI3K-Akt and JAK-STAT signalling (Figs. 4B, 5B, 6B, Table S2). Gene ontology (GO) analysis of target genes in these networks identified enrichment of genes related to ‘myelin sheath’ across the three networks (Table S3). Specific miRNA targets of myelination genes included *Padi2* (regulated by miR-3473b) and *Hmgcr* (regulated miR-29a-3p). In addition, pathways related to neurodegenerative disease were overrepresented in all networks. These age-specific regulatory networks were primarily composed of miRNAs that were enriched in all age groups, and therefore define robust regulation of conserved microglial transcriptional networks during CNS development and adult homeostasis.

**Figure 4.**
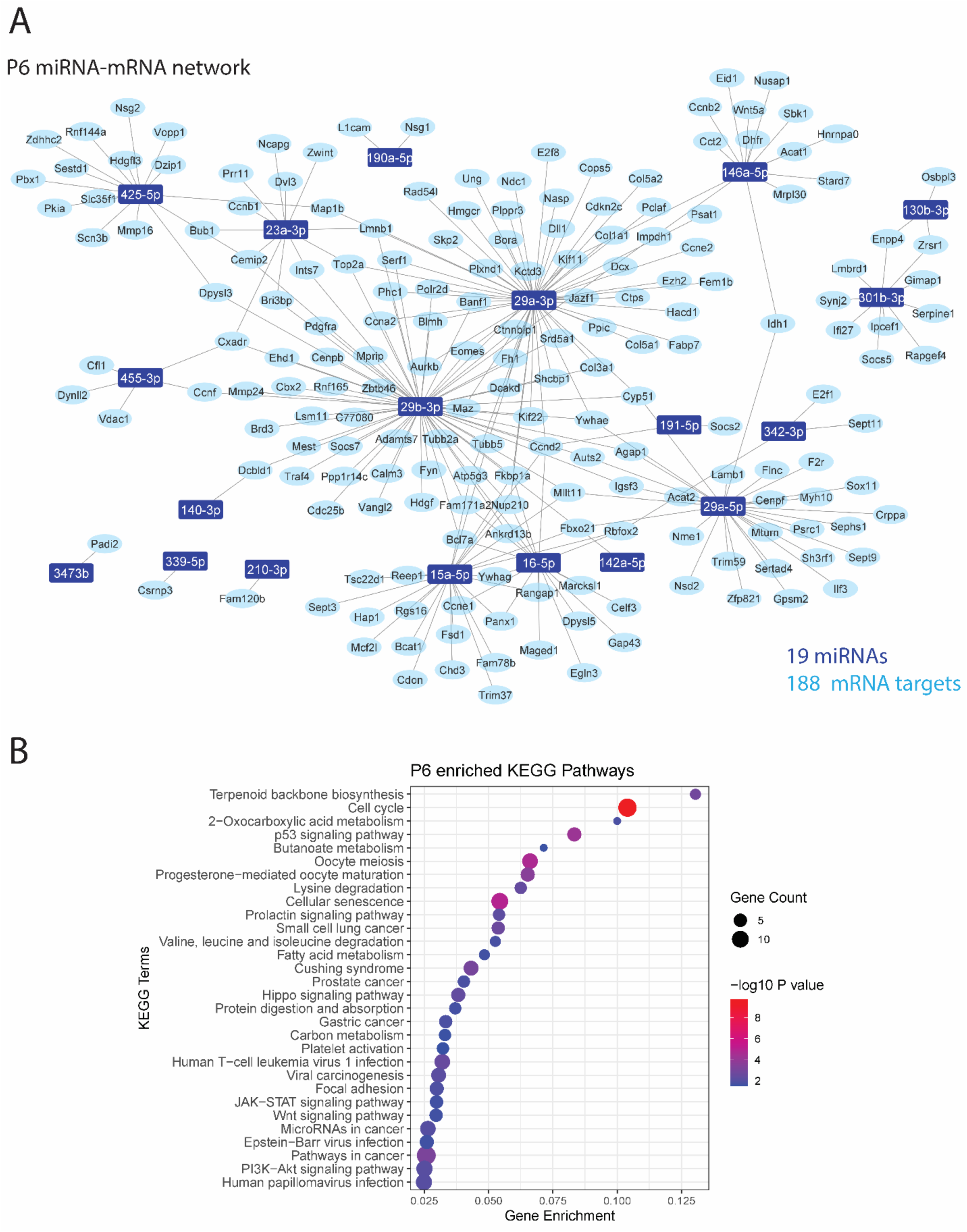
miRNA-mRNA coregulatory network highlight important CNS processes regulated by P6 microglia. **A**. miRNA-mRNA regulatory networks were constructed from negative correlations (r < -0.857, FDR < 0.05) between miRNA and mRNA expression data generated from P6 microglia and validated with external evidence for interaction. **B**. KEGG enrichment dot plots were generated from target genes in the network. Top 30 pathways/terms (by P value) are represented in each plot and ordered by Gene Enrichment score (number of genes in the dataset represented in the gene set/total number of genes in the gene set).

**Figure 5.**
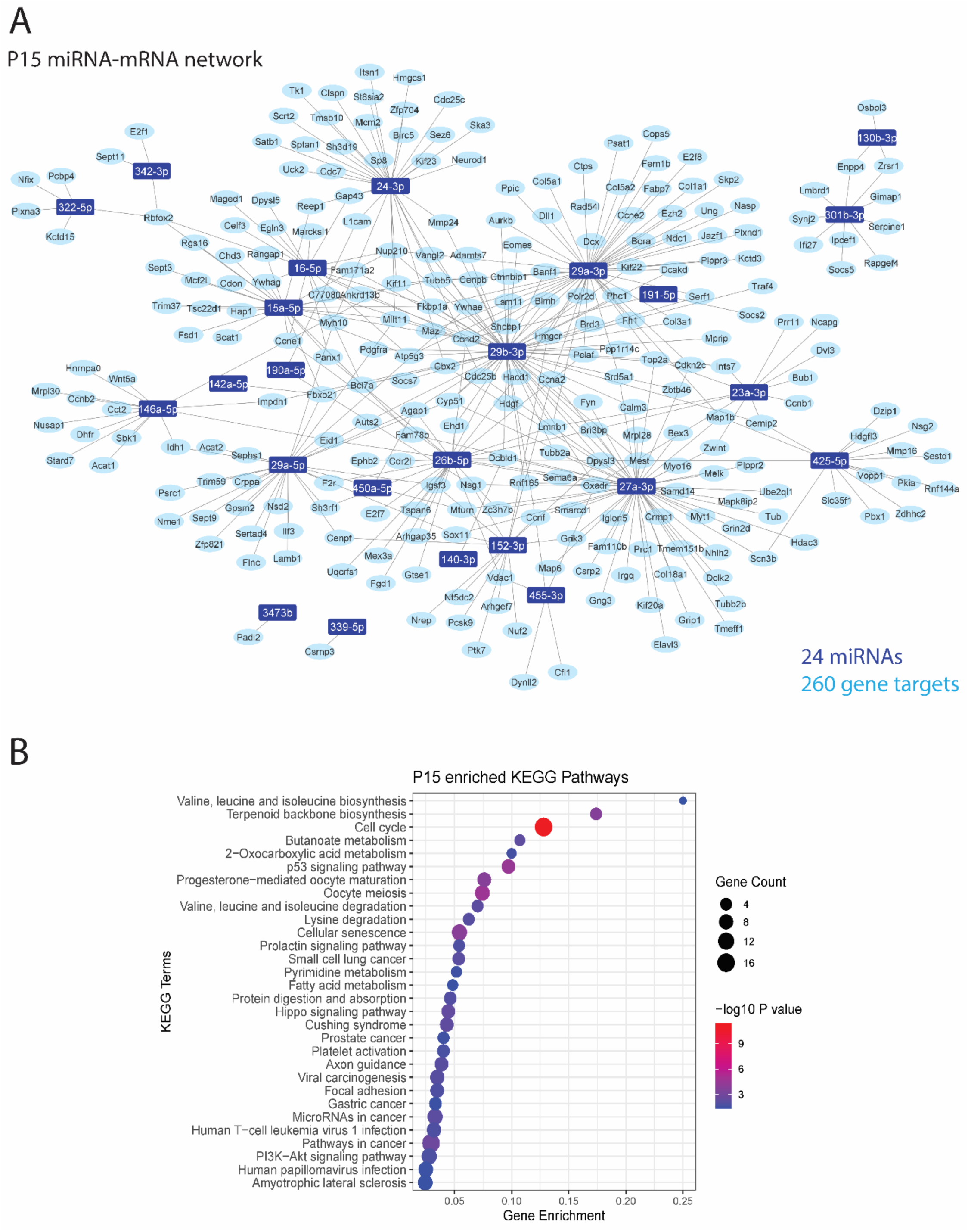
miRNA-mRNA coregulatory network highlight important CNS processes regulated by P15 microglia. **A**. miRNA-mRNA regulatory networks were constructed from negative correlations (r < -0.857, FDR < 0.05) between miRNA and mRNA expression data generated from P15 microglia and validated with external evidence for interaction. **B**. KEGG enrichment dot plots were generated from target genes in the network. Top 30 pathways/terms (by P value) are represented in each plot and ordered by Gene enrichment score (number of genes in the dataset represented in the gene set/total number of genes in the gene set).

**Figure 6.**
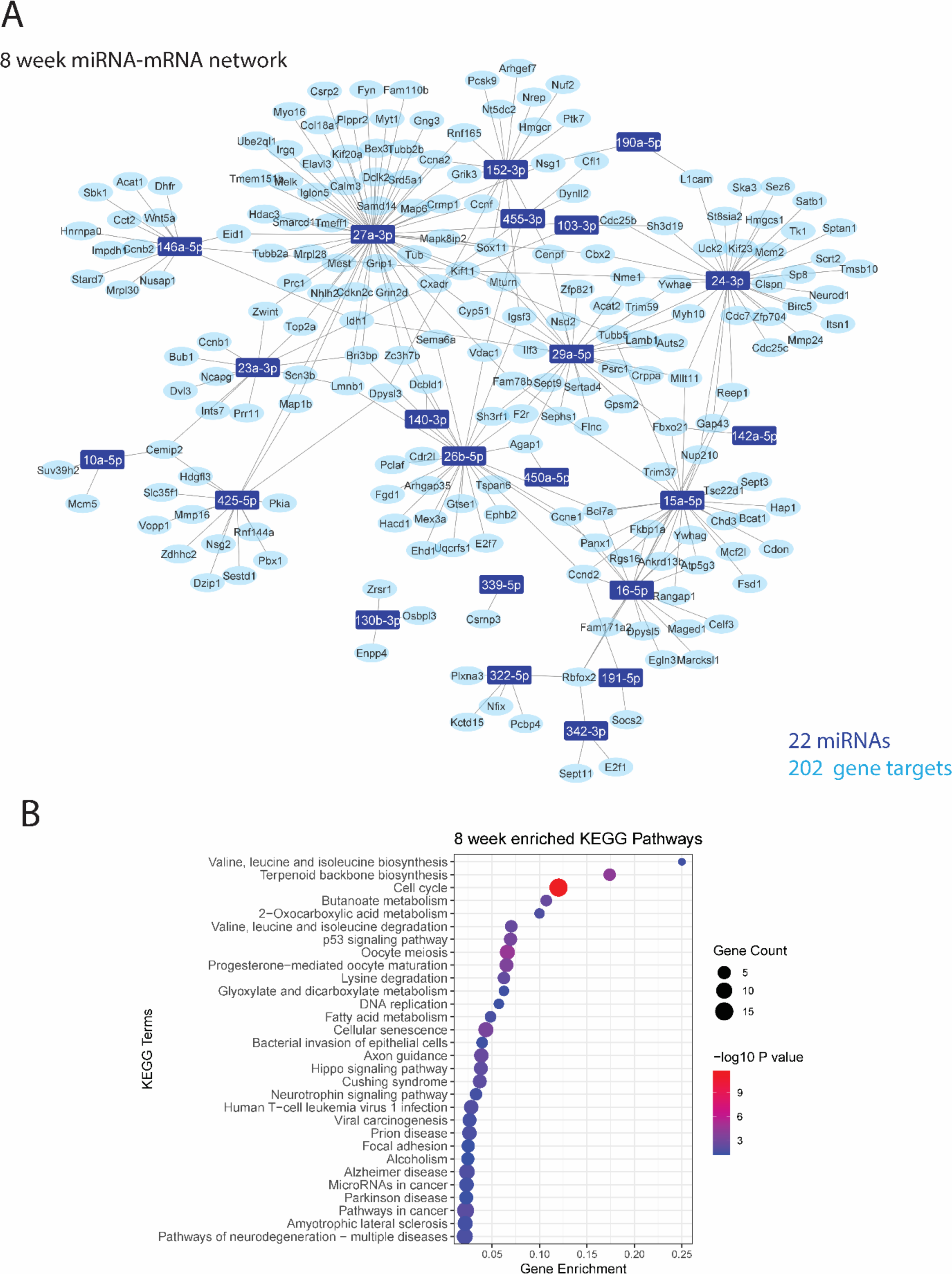
miRNA-mRNA coregulatory network highlight important CNS processes regulated by 8 week microglia. **A**. miRNA-mRNA regulatory networks were constructed from negative correlations (r < -0.857, FDR < 0.05) between miRNA and mRNA expression data generated from 8 week microglia and validated with external evidence for interaction. **B**. KEGG enrichment dot plots were generated from target genes in the network. Top 30 pathways/terms (by P value) are represented in each plot and ordered by Gene enrichment score (number of genes in the dataset represented in the gene set/total number of genes in the gene set).

### Subsets of temporally dynamic miRNAs regulate specific aspects of microglial development

Microglia undergo major phenotypic and transcriptional transitions from early post-natal development through to adulthood^21^. In order to identify the miRNAs that are most likely to regulate developmental transitions in microglia, we next interrogated changes in miRNA expression between age groups. We performed direct pairwise comparisons of miRNA expression profiles between different development ages to define subsets of differentially expressed miRNAs (Fig. 7A, Table S4). Some 140 differentially expressed miRNAs were observed across all comparisons, with the largest difference observed between P6 and 8-week microglia, reflecting the significant maturation of developmental microglia to their adult homeostatic counterparts (Fig. 7B). To further characterise the individual developmental trajectories of differentially expressed miRNAs, we plotted fold changes of expression at P15 and 8 weeks, relative to P6 and categorised significantly altered miRNAs into groups based on their changing patterns of expression (Fig. 7C). This analysis identified 19 miRNAs that were sequentially upregulated, and 35 miRNAs that were sequentially downregulated between P6 and adult microglia (Fig. 7C and 7D). Gene ontology (GO) analysis of the predicted target genes for the upregulated miRNA subset were strongly related to cyclin activity and cell cycle transition consistent with the proliferative nature of early post-natal microglia^31^(Fig. 7E). In contrast, predicted gene targets of the downregulated miRNA subset were overrepresented in biological processes related to wound healing, endocytosis, migration and immune function (Fig. 7E).

**Figure 7:**
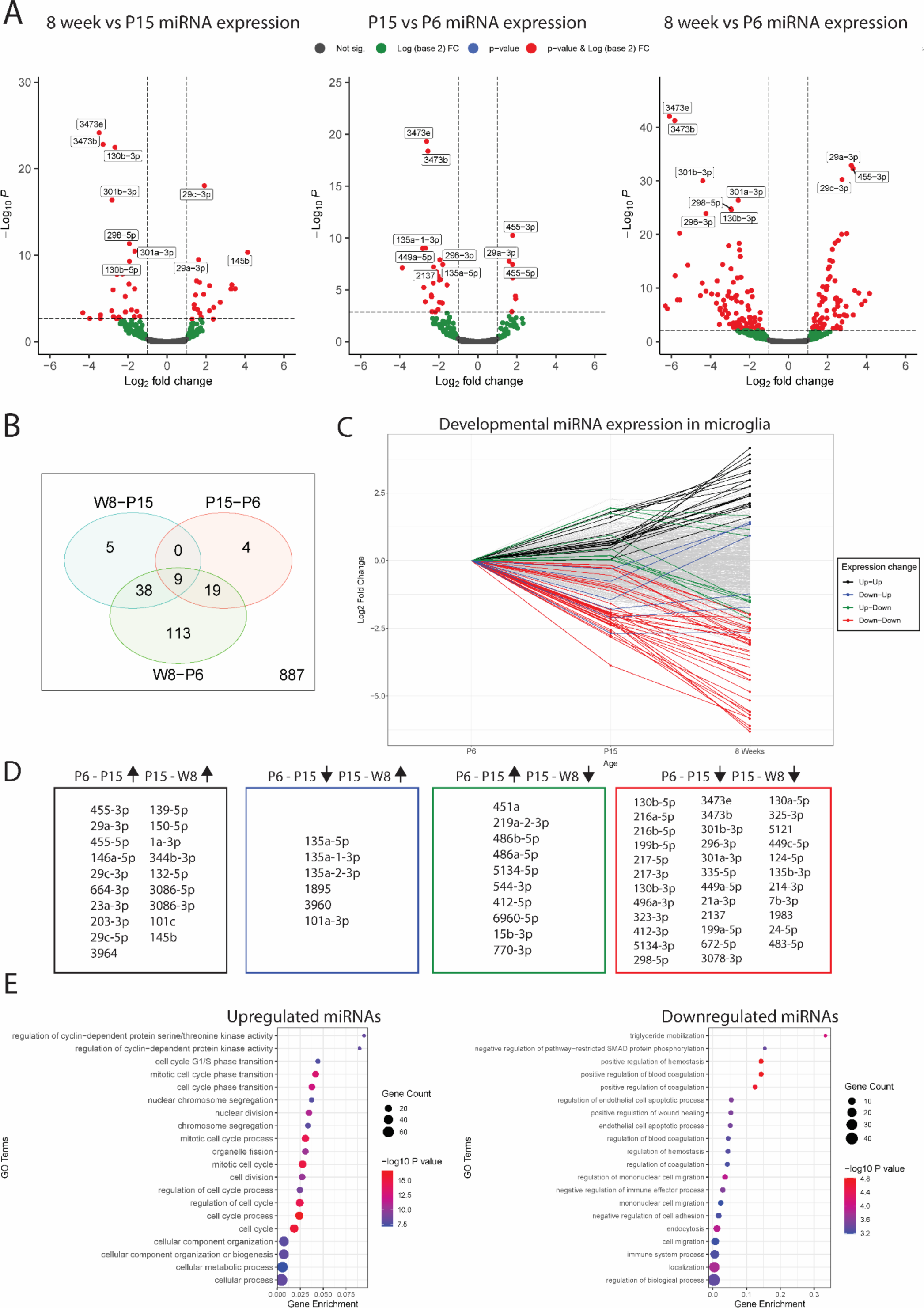
Subsets of microglial miRNAs are differentially expressed over developmental time. **A**. Volcano plots of differentially expressed miRNAs of three pairwise comparisons of P6, P15 and 8 week (adult) microglia populations. Top 10 miRNAs (by p value) are labelled in each plot. **B**. Venn diagrams comparing differentially expressed (logFC >1, FDR < 0.05) miRNAs from pairwise comparisons of age groups. 140 miRNAs were differentially expressed in at least one comparison. **C** miRNA expression (n=1075) was plotted as LogFC relative to expression at P6 to observe changes over developmental time. miRNAs with at least one significant fold change between age group transitions (P6-P15 and/or P15-W8) were categorised by their pattern of expression (ie. up-up, up-down, down-up or down-down). Presence of a point at a specific age group indicates that specific transition as statistically significant (FDR < 0.05). **D**. Lists of miRNAs corresponding to each category as described in 5C. The largest subsets were the up-up and down-down categories, with 19 and 35 miRNAs respectively. **E**. Gene Ontology (GO) analysis of predicted target genes of consistently upregulated (up-up) and downregulated (down-down) miRNAs (related to Fig 5A). Top 20 pathways/terms (by P value) are represented in each plot and ordered by Gene enrichment score (number of genes in the dataset represented in the specific gene set/total number of genes in the specific gene set).

### Analysis of novel miRNAs enriched in microglia identifies chr3_29427 as a putative novel regulator of microglial biology

Of the 1075 miRNAs detected across all samples, 53 were identified as previously unannotated novel miRNA sequences by miRDeep2^32^. Twenty six of these novel sequences were specifically enriched in microglia relative to CD45^-ve^ CNS cells (Fig. 8A; Table 3). The most strongly expressed novel miRNA candidate in this regard was chr3_29427, which mapped to an intergenic region on chromosome 3 (chr3:150,866,217 - 150,866,276, GRCm38/mm10) and was enriched at a minimum of 16-fold across all ages and observed to increase with expression with age (Table 3). We validated its strong enrichment (but not increasing expression with age) by qPCR (Fig. 8B). Next, we constructed novel miRNA-mRNA regulatory networks, filtering for strongly negative correlations between miRNA-mRNA pairs (r < -0.895, FDR < 0.05). In the absence of experimental evidence for novel miRNA binding interactions, we calculated predicted interactions using the hybridization tool RNAhybrid, filtering for those that were strongly predicted to bind with a minimum free fold energy < 25 kcal/mol (p < 0.05)^33^. Twenty-six miRNA-mRNA interactions passed these criteria, all of which included the top novel miRNA candidate chr3_29427 (Fig 8C). Linear modelling plotting chr3_29427 expression against expression of three predicted targets in this regulatory network; Fam171a2, Birc5 and Cdca5 highlights a consistent pattern of increasing chr3_29427 expression associated with downregulation of target genes to a negligible level of expression in 8 week microglia (Fig. 8D).

**Table 3.**
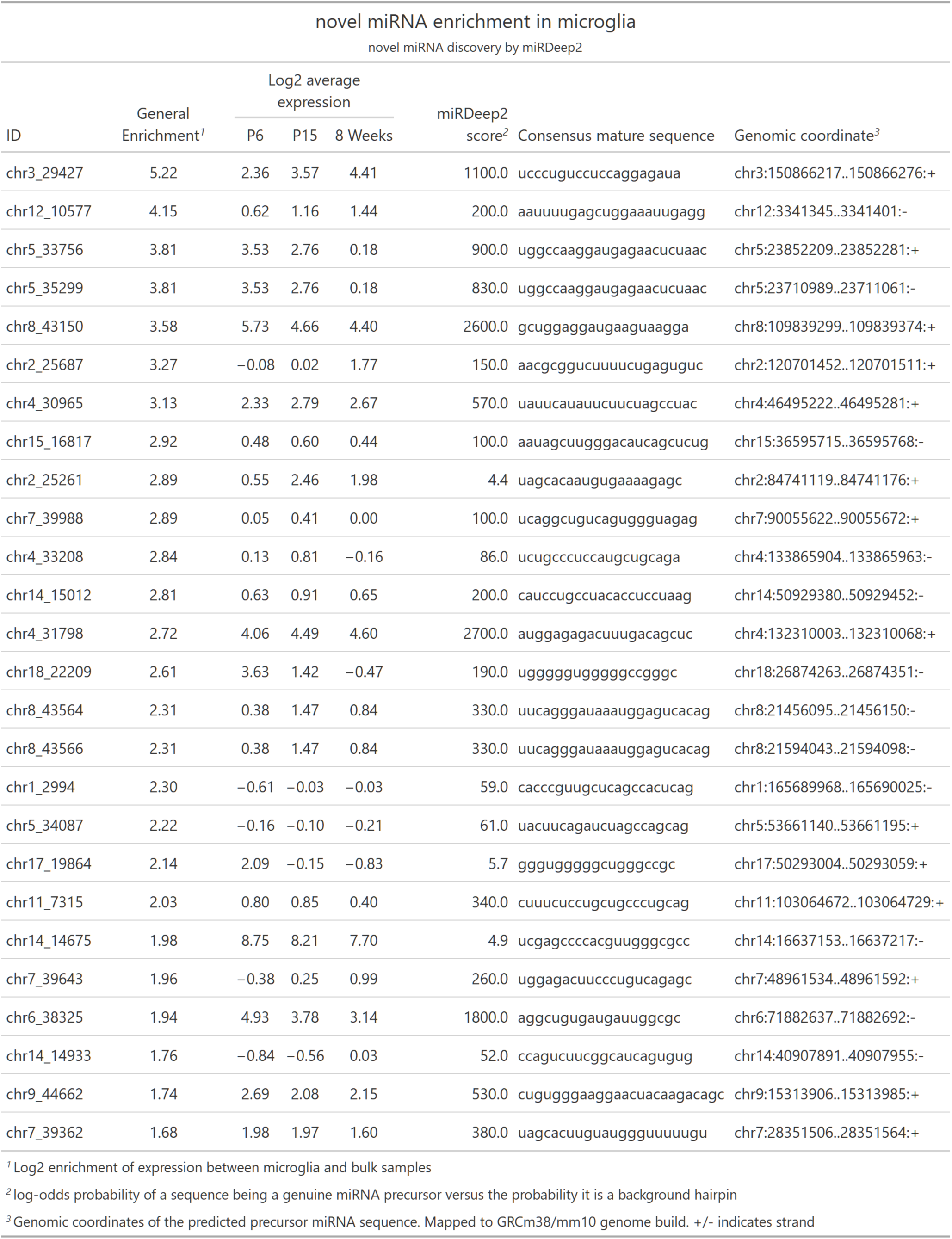

**Figure 8.**
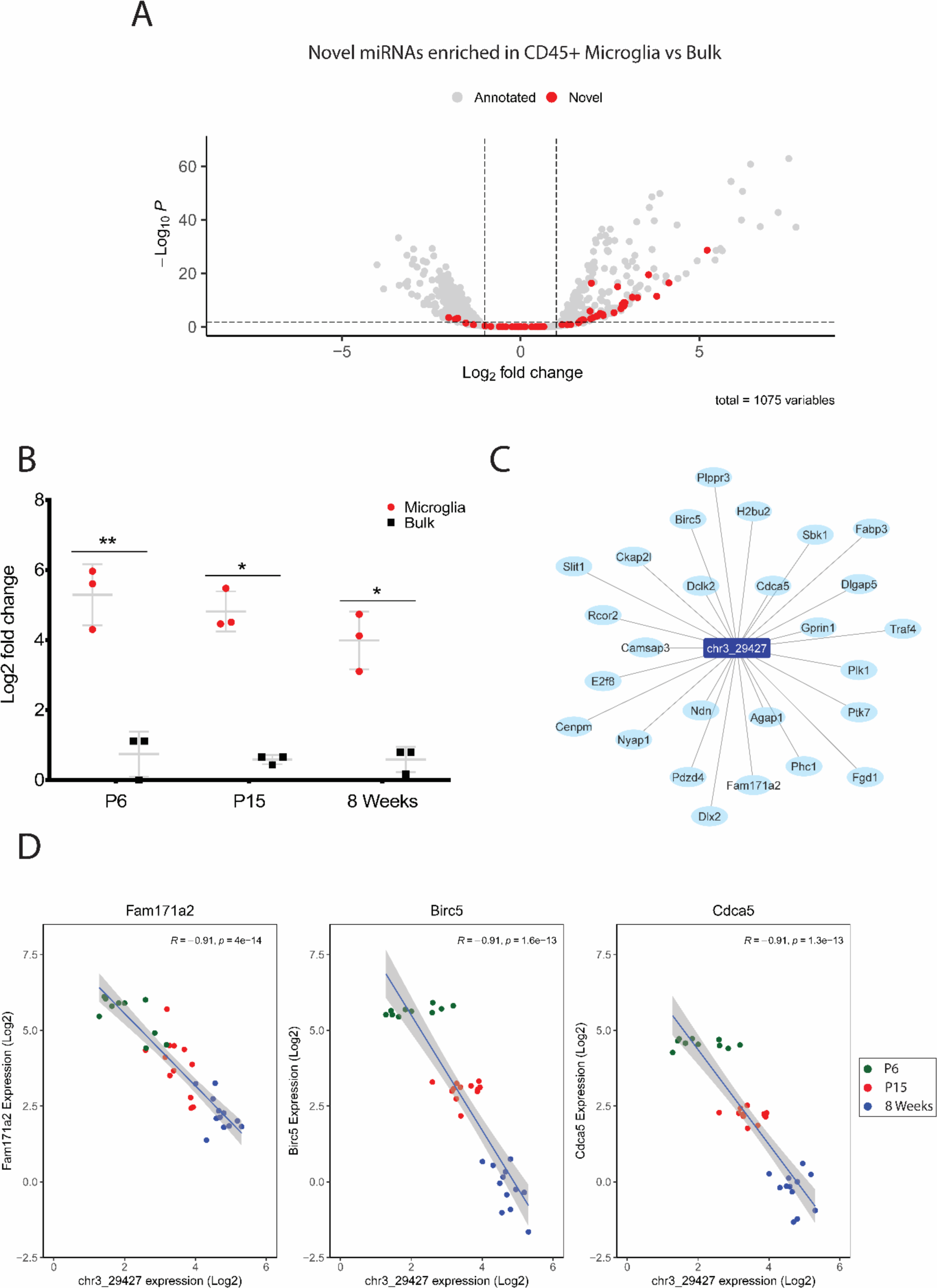
miRNA discovery pipeline identified 26 novel miRNA candidates enriched in developmental microglia. **A**. Volcano plot of differential miRNA expression between pooled microglia and bulk samples, labelled as either novel (red) or annotated (grey) (related to Fig 2A). Of the 56 novel miRNAs detected in all samples, 26 were identified as generally enriched in microglia. **B**. qPCR expression analysis of chr3_29427 in microglia and bulk samples across developmental time (n = 3 males in each group). Significant enrichment was observed at all ages. Data were analysed by paired t tests comparing chr3_29427 expression between microglia and bulk in each age group. * p< 0.05, ** p < 0.01. Lines on qPCR plots represent mean and standard deviation. **C**. chr3_29427-mRNA regulatory network constructed from strong negative correlations (r < -0.857, FDR < 0.05) from mRNA expression data and prediction of interaction using RNAHybrid. 26 putative targets were identified for chr3_29427. **D**. Log2 expression of chr3_295427 and 3 of its putative targets; Fam171a2, Birc5 and Cdca5. Linear modelling identifies strong negative relationships between the novel candidate and all 3 targets.

### Male and female developmental microglia exhibit no differential miRNA expression and a small difference in mRNA expression

Mouse cohorts in this study were balanced for sex, based on previous reports of sexual dimorphism in microglial genetics^34–36^. However, comparisons of miRNA and mRNA expression between sexes within each age group revealed no differentially expressed miRNAs and only 23 differentially expressed mRNAs (Fig. 9A and 9B). To further validate the absence of sexually dimorphic miRNAs, we used qPCR to directly assess the expression of candidate miRNAs previously identified as sexually dimorphic in adult microglia^35^. None of the selected candidates exhibited differential expression between sexes in 8 week old mice. (Fig. 9C). Conversely, mRNAs which were differentially expressed between sexes were observed in each age group, with 14 being specific to 8 week mice (Fig. 9B, Fig S2). Amongst genes differentially expressed at all ages were *Xist* and *Ddx3y*, previously identified as specifically expressed in female and male microglia respectively^20,37^. For P6 and P15 mice, every sex-specific gene, except for *Il10* in P15 mice, was located on the X or Y chromosome (Table 4). Seven of the 22 sexually dimorphic genes observed in adult mice were autosomal. Amongst the genes that were identified as sexually dimorphic were several related to microglial immune function, including *Cd44, Cxcr2, S100a8, S100a9* and *Il10*^38–41^ (Table 4). *S100a8* and *S100a9* have been previously identified as upregulated in male microglia isolated from adult mice^34,42^.

**Table 4:**

| Differential gene expression between male and female microglia |  |  |  |  |  |  |  |
| --- | --- | --- | --- | --- | --- | --- | --- |
| Sexually dimorphic gene expression in P6, P15, and 8 week microglia |  |  |  |  |  |  |  |
| Gene | LogFC<sup>1</sup> | Adjusted p value<sup>2</sup> | Chromosome | Gene | LogFC<sup>1</sup> | Adjusted p value<sup>2</sup> | Chromosome |
| 8 weeks |  |  |  | P15 |  |  |  |
| Kdm6a | -1.2776291 | 4.684453e-03 | X | Il10 | 1.9426059 | 2.513089e-02 | 1 |
| Xist | -10.8392421 | 3.417637e-16 | X | Xist | -9.9889926 | 5.855535e-17 | X |
| Kdm5d | 10.7478933 | 3.800953e-11 | Y | Kdm5d | 9.5232972 | 1.056057e-09 | Y |
| Tsix | -4.4852594 | 1.561152e-08 | X | Tsix | -4.9033329 | 1.457855e-09 | X |
| Uty | 9.2863139 | 7.767534e-08 | Y | Uty | 8.7677246 | 3.426706e-07 | Y |
| Ddx3y | 10.3057299 | 7.420489e-06 | Y | Eif2s3x | -0.9007106 | 3.312759e-05 | X |
| Eif2s3y | 10.4986331 | 5.789245e-05 | Y | Ddx3y | 9.2600469 | 4.737304e-05 | Y |
| Bst1 | 1.7110444 | 1.503425e-03 | 5 | Eif2s3y | 9.1902288 | 2.856358e-04 | Y |
| Cxcr2 | 1.4639865 | 2.306832e-03 | 1 | P6 |  |  |  |
| Cd44 | 0.8368712 | 2.754180e-03 | 2 | Xist | -10.8213305 | 5.266675e-15 | X |
| Adgre4 | 1.2744018 | 7.172776e-03 | 17 | Kdm5d | 10.4343516 | 1.595254e-08 | Y |
| S100a8 | 1.7293343 | 1.016865e-02 | 3 | Tsix | -4.5835985 | 3.454569e-08 | X |
| Mefv | 1.9139963 | 1.380656e-02 | 16 | Uty | 9.8287674 | 2.184195e-06 | Y |
| S100a9 | 1.6754487 | 1.528078e-02 | 3 | Ddx3y | 10.6196418 | 1.112405e-03 | Y |
| Hp | 1.5682624 | 1.661822e-02 | 8 | Eif2s3y | 10.7382174 | 1.766455e-03 | Y |
| Slfn1 | 1.1997261 | 1.992361e-02 | 11 |  |  |  |  |
| Kdm5c | -0.4240698 | 2.748662e-02 | X |  |  |  |  |
| Eif2s3x | -0.5565456 | 3.482553e-02 | X |  |  |  |  |
| Emilin2 | 1.1161409 | 3.482553e-02 | 17 |  |  |  |  |
| Lrg1 | 2.2529933 | 3.482553e-02 | 17 |  |  |  |  |
| Ifitm2 | 0.8528811 | 3.482553e-02 | 7 |  |  |  |  |
| Slfn4 | 1.5351021 | 4.382920e-02 | 11 |  |  |  |  |
| <sup>1</sup> LogFC = Log2 fold change comparing male to female gene expression |  |  |  |  |  |  |  |
| <sup>2</sup> FDR adjusted |  |  |  |  |  |  |  |

**Figure 9:**
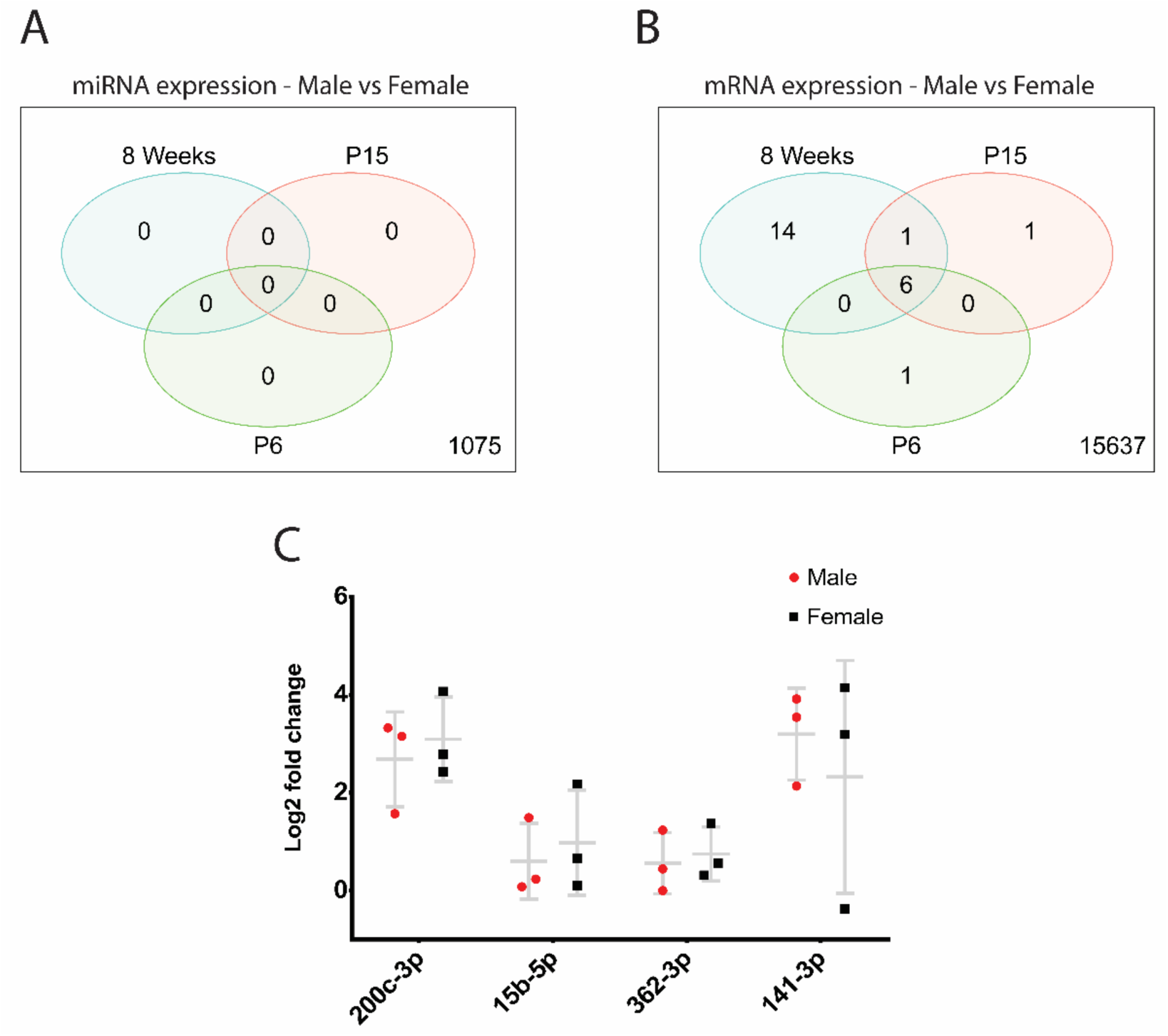
Differential miRNA and mRNA expression between male and female microglia across developmental age. **A**. Venn diagram comparing differential expression of miRNAs between male and female microglia in each age group. No miRNAs were differentially expressed at any age (FDR <0.05). **B**. Venn diagram comparing differential expression of mRNA between male and female microglia in each age group. 23 mRNAs were identified as differentially expressed in at least one age group, with most of the differential expression observed in 8 week mice. A full list of differentially expressed genes is outlined in Table 2. **C**. qPCR expression of selected miRNA candidates between male and female microglia (n=3 in each group). Data were analysed by unpaired t tests comparing microglia and bulk group expression for each tested miRNA. No statistically significant differences in expression were observed for any of the tested miRNAs.

## Discussion

In this study we have characterised the miRNAome of developmental microglia, identifying both a core miRNA signature enriched in microglia, as well as unique age-specific subsets of miRNAs. We have constructed a miRNA-mRNA network for microglia, which implicate both previously annotated and novel miRNAs in the regulation of key neurodevelopmental processes, including neurogenesis and myelination. Unexpectedly, we did not observe sex-specific miRNA expression, along with only a handful of sex-specific mRNA transcripts, indicating that during developmental microglia are not necessarily inherently different between sexes. Our developmentally focused dataset provide insight into the epigenetic mechanisms regulating the transition of microglia from a proliferative and highly reactive cell population to a mature homeostatic population within the CNS.

We identified a strong miRNA enrichment signature in microglia when compared to the surrounding CNS cells, with over 300 miRNAs upregulated in microglia in at least one age group. This was not unexpected, as studies of microglial transcriptomes have highlighted these cells as having a unique transcriptional signature^43–45^. Unlike the other major CNS cell types (astrocytes, neurons, oligodendrocytes), microglia do not derive from neuroepithelial lineages, but instead arise from the embryonic yolk sac before migrating to and establishing a self-sufficient population in the developing CNS^46–48^. Our microglial mRNA data also reflected this distinct signature. Amongst the top expressed mRNA transcripts in this sequencing data were well-defined microglial identity genes including *Cx3cr1, Csf1r, Sparc, Tmem119* and *Hexb*^49^. Together, these results highlight a unique genetic identity of microglia compared to the surrounding CNS environment.

The present study has defined a dynamic microglial miRNA expression profile, comprised of a ‘core’ signature of consistently enriched miRNAs and subsets of temporally specific miRNAs. In conjunction with integrated miRNA-mRNA network analysis, we were able to characterise age-specific domains of miRNA expression and explore their influence upon microglial mRNA transcription throughout development. Firstly, 91 miRNAs were observed to be significantly enriched in microglia at all ages compared to other CNS cell types. Many of the highly expressed miRNAs in this subset have been previously identified in microglia, including miR-146, miR-223 and miR-142^23,50^. These miRNAs have defined functions in the homeostatic regulation of microglial phenotype and their dysregulation is linked to inflammation and neurodegeneration^34,51–54^. In addition to this common microglial signature, this study identified 129 miRNAs that were specifically enriched in a single age group. Although most of these miRNAs were expressed in adult cells there was significant enrichment of specific miRNAs at all ages, suggesting that miRNAs could be important for regulating age-specific microglial functions. This was further supported by correlated miRNA-mRNA expression, which implicated specific gene pathways as targets of miRNA regulation in developmental microglia.

To fully characterise the roles of miRNAs in cellular systems it is important to define the gene networks which they regulate. miRNA-mRNA networks in this study were constructed using both stringent correlation thresholds and experimental validation of miRNA-mRNA interaction. Pathway analysis of target mRNAs in age-specific networks were strongly enriched for cell cycle related processes. Developmental microglia proliferate as they colonise the brain. This includes a significant wave of proliferation that peaks during the second post-natal week of development, before numbers stabilise in adulthood^31^. Ablation and perturbation of the developmental microglial population can have long lasting effects on the integrity of neurons and therefore the regulation of microglial proliferation is essential for normal CNS development^55^. Network analysis identified numerous miRNA candidates that could provide important regulatory influence on the microglial cell cycle. For example, the miR-301/130b cluster was identified as strongly enriched in P6 and P15 microglia. This cluster is associated with hematopoietic cell proliferation and migration and identified targets of this cluster in microglia in our data included *Serpine1, Socs5* and *Ifi27* which are implicated in cell proliferation, motility, interferon signalling and endocytosis^56–60^. Also enriched in microglia was the miR-29 cluster (miR-29a, miR-29b), which is also associated with cell proliferation dynamics and downregulation of cyclins and cyclin dependent kinases^57,61^. Another interesting candidate for the regulation of microglial development is miR-10a-5p. Previously uncharacterised in microglia, miR-10a-5p, is uniquely enriched in adult mice and targets and therefore potentially inhibits Mcm5, an established regulator of microglial cell cycle that is strongly expressed during embryonic and early post-natal microglial development^21,43^.

A central role for developmental miRNAs as regulators of cell cycle progression in microglia was reinforced by comparisons of disparate miRNA expression between microglia isolated from P6 and 8 week mice. This analysis identified 19 miRNAs that were consistently upregulated between P6 and 8 weeks, and whose putative gene targets were overwhelmingly associated with cell cycle. This included the members of the miR-29 family; miR-29a-3p and miR-29c-3p. Interestingly, expression of the miR-29 progressively increases in ageing microglia and is associated with acquisition of an inflammatory microglial phenotype, indicating that it may regulate several processes in microglia^62^. Our data suggests that subsets of miRNAs in microglia target key genes associated with cell cycle progression and inhibiting proliferation as developing microglia transition to homeostasis.

In addition to cell cycle regulation, pathway analysis of miRNA-mRNA networks revealed an overrepresentation of processes related to CNS development, including neurogenesis, axon guidance and myelination. Microglia are essential modulators of these processes and their dysregulation is a major contributor towards neurodevelopmental disorders^63^. Regulation of neurogenesis by microglia can occur both directly, via efferocytosis of either apoptotic immature neurons, synapses or oligodendrocytes and indirectly via secretion of neurotrophic factors including IGF-1 and BDNF to support neural cell growth and survival^64^. Amongst neurogenesis related genes represented in miRNA-mRNA networks were Dypls3 (targeted by miR-425-3p and miR-29b-3p) and Dypls5 (targeted by miR-16b-5p), cytoskeleton associated genes that are strongly enriched in ameboid microglia during the first week of postnatal development^7,65,66^. Knockdown of Dypls3 inhibits microglial phagocytosis and migration and therefore could be important for maintaining a specific microglial phenotype associated with post-natal neurogenesis^65^. Microglia also regulate developmental myelination and maintain white matter integrity. In addition to clearance of immature oligodendrocytes and myelin debris, microglia secrete trophic factors to support OPC development, lipid metabolism and myelination^2,3,67^. We identified several interesting miRNA-mRNA interactions that may mediate these processes. This included miR-29a-3p targeting Hmgcr, a regulator of cholesterol homeostasis that is enriched in white matter associated microglia during myelinogenesis^68^. Another interesting candidate is miR-3473b, represented in both the P6 and P15 networks that targets a single gene, *PADI2*. PADI2 regulates citrullination of myelin basic protein (MBP), which alters myelin ultrastructure. Of note, it has been hypothesised that myelin modifications of this nature could be a source of autoimmune pathogens potentially implicated in the pathogenesis of MS^69^. PADI2 is enriched in subsets of microglia isolated from human MS tissue and therefore miR-3473b could play an important role in refining PADI2 expression, both during development and in demyelinating contexts^19^.

Post-natal microglia are active participants in apoptotic clearance of neural cells within the developing CNS, where they adopt a reactive and highly phagocytic ameboid phenotype^31^. In the transition to adulthood, microglia shift towards a ramified phenotype concordant with a stronger emphasis on immune surveillance. In conjunction with identifying a subset of 19 temporally upregulated miRNAs that regulate cell cycle, temporal analysis of miRNA expression also revealed a subset of 35 miRNAs that were strongly downregulated with increasing age. KEGG analysis of the mRNA targets of this downregulated subset indicated that these miRNAs were associated with processes related to wound healing, mononuclear cell migration and immune function^31^. The downregulation of these miRNAs in 8-week microglia may therefore be associated with the upregulation of immune functions that are predominant in adult cells and associated with homeostasis. This forms an interesting contrast with the cell-cycle related miRNAs that were upregulated, whereby these miRNA subsets may be essential in co-ordinating microglial transition to adult homeostasis, simultaneously suppressing cell proliferation and upregulating immune effector functions.

More broadly, pathway analysis of targets in all miRNA-mRNA networks identified a significant enrichment of pathways related to inflammatory signaling pathways and immune activation. This included PI3k-Akt and JAK-STAT signaling, central regulators of microglial mediated neuroinflammation^70,71^. We also observed enrichment of miRNAs with links to inflammation and neurodegeneration which are not well defined in microglia including miR-511-3p, miR-1895 and miR-6983-5p. miR-511-3p has been characterised in tumor and lung associated macrophage populations, encoded by and co-regulated with the CD206 gene, where it is associated with anti-inflammatory (M2-like) activation^72,73^. miR-6983 is downregulated in oligodendrocytes experimental autoimmune encephalomyelitis (EAE), an animal model of demyelination, and is associated with disrupted ferroptosis pathways that are implicated in demyelination and neurodegeneration^74^. Interestingly, miR-210-3p, which was uniquely enriched in the P6 miRNA network, is associated with neurotoxic microglial activation in a P7 mouse model of neo-natal ischemic encephalopathy, making it a potentially unique modulator of early developmental inflammation^75^. With functions related to macrophage polarisation and pathways of neurodegeneration, further work is required to elucidate specific roles for these highly expressed miRNAs in microglia.

The stringent network analysis method employed in this study refined networks such that only high confidence interactions between miRNAs and mRNAs were included, increasing the robustness of predicted genetic networks. There was a significant overlap of represented miRNAs when comparing across age networks, indicating that they were primarily composed of miRNAs with conserved enrichment throughout development. Of the 129 miRNAs which were specifically enriched at a single developmental time point, only 3 were represented in these networks: miR-210-3p (enriched at P6), miR-10a-5p (enriched at 8 weeks) and miR-103-3p (enriched at 8 weeks). As discussed above, these uniquely enriched miRNAs are implicated in important microglial processes. By employing such a stringent correlation threshold, this study maximised the predictive potential of miRNA-mRNA interactions at the cost of potentially missing other miRNA-mRNA interactions, including those that are specific to a developmental time point.

In addition to previously annotated miRNAs, we identified 26 novel miRNA candidates enriched in microglia. Many of these, including the highly expressed chr3_29427, may regulate important microglial functions in the CNS. Amongst putative mRNA targets were key regulators of cell cycle regulation and neurogenesis. These included Fam171A2, which regulates progranulin expression and modifies risk of neurodegeneration^76^. Cell cycle genes Cdca5 and Birc5 were also identified as putative targets of chr3_2942. Birc5 has been previously characterised as highly expressed by proliferating microglia proliferation during development and in response to injury^77^. These predicted novel candidates may be critical regulators of unique microglial biology and warrant future functional validation *in vitro* and *in vivo*.

We did not identify any association of sex with microglial miRNA abundance. Specifically, 23 mRNAs but no miRNAs were identified as differentially expressed between male and female microglia across development. There is growing evidence to suggest that sex hormones significantly influence the developmental trajectories of male and female microglia manifesting in differential regional densities, morphology, phagocytic capacity and metabolic influx^78^. Importantly, these innate differences are exacerbated during immune challenge and stress and are suggested to be a prominent influence in observed sex biases of neurological disease^34,36,42,79^. Transcriptomics studies have defined subsets of mRNAs and miRNAs that are differentially expressed between male and female microglia, in steady state and particularly following immune challenge in adult mice^34,35,42,79,80^. In contrast, we did not observe any differentially expressed miRNAs, a finding which was further corroborated with qPCR analysis of four candidates that were previously identified as sexually dimorphic at baseline in 6 month old B6C3F1 mice^35^. Conversely, our study identified 23 differentially expressed mRNA transcripts across development, with the majority of these being located on sex chromosomes and specifically differentiated in adult mice. The robust and predictable differentiation of sex-linked genes including Xist and Ddx3y support the veracity of the dataset. Additionally, male specific enrichment of S100a9 has been observed in both our dataset and in related studies^34,42^. We did identify a small subset of immune related genes as significantly upregulated in male microglia including, Cxcr2, Il10 and CD44 all of which are dysregulated during inflammatory microglial activation^39,41,81^. Overall, this study has identified far fewer differentially expressed genes than related scientific studies.

What might be causing this discrepancy? Interestingly, while functional and structural differences between male and female microglia have been consistently observed, transcriptomic results are less congruent^78^. Firstly, age seems to be an important factor. A comprehensive developmental study by Hanamsagar et al. observed that transcriptionally, male and female microglia are quite similar until early adulthood (P60). Other reported significant transcriptional differences studies come mainly from adult mice, (12 weeks and older) reinforcing the idea that the effects of sex are more prevalent with age^34,42,80^. However, our dataset and others have reported a minimal effect of sex even in young adult cells (8 weeks) ^16,20^. Further studies that assess transcription at expanded time points (beyond 8 weeks of development) will assist in an accurate representation of the effect of sexual development on microglial gene expression. Another hypothesis for these discrepancies is the method of microglial isolation. Microglial cells are dynamic and respond quickly to changes in their local environment^15^. While an unavoidable caveat, it has been shown that different methods of isolation (ie. immunopanning, FACS, magnetic bead sorting) can differentially alter the activation status of microglia and hence gene expression, potentially contributing to differing reports of baseline sexual dimorphism in studies of microglia^45^.

Additionally, aspects of sexual dimorphism in microglial gene expression are region dependent^36,42^. Sexually dimorphic region specific signatures would be diluted in the current study, given that the microglial populations were derived from the whole brain. Nevertheless, this result adds another perspective to the growing discussion of sexual dimorphism of microglia and suggests that sex may have a lesser influence on microglial transcription than previously reported. Specifically, dimorphism may be more prevalent post-activation, whether induced by isolation, immune challenge or in ageing related neurodegeneration.

In summary, we characterised a unique and highly controlled dataset of miRNA expression in developing microglia, capturing a dynamic profile that may contribute to the complex roles that microglia adopt in post-natal development and the establishment of homeostasis. This miRNAome is composed of both a relatively unchanging subset of highly enriched miRNAs and subsets of miRNAs that exhibit age-dependent expression profiles. Correlation of known and novel miRNAs with mRNA expression data enabled the identification of target mRNA networks that mediate neurodevelopmental and immune processes that are central to microglial function in the CNS. In combination with environmental cues, epigenetic mechanisms shape the genetic landscape of microglia and refine phenotypes. Dysregulation of these epigenetic mechanisms can be a major driver of neurodevelopmental and neurodegenerative disorders. This developmental profile of paired miRNA and mRNA expression represents an important step in understanding the genetic regulation of microglia and their impact on the CNS in health and disease.

## Materials and methods

### Experimental and analytical design

The experimental paradigm was designed to identify miRNAs expressed specifically in microglia compared with other CNS cells across three developmental timepoints; P6, P15 and 8 weeks. In addition, all groups contained an equal number of male and female mice to allow for identification of any sex specific miRNAs. An overview of the experimental and analytical design is given in figure 1.

### Chemicals and reagents

All chemicals were purchased from Sigma (CA, USA) and all cell culture media were purchased from Gibco (TX, USA) unless otherwise stated.

### Isolation of microglial populations from murine brain tissue

All animal experiments were approved by the animal ethics committee at the Florey Institute of Neuroscience and Mental Health and conducted in accordance with the National Health and Medical Research Council Guidelines. Whole brain was extracted from mice at appropriate ages (Fig 1). Microglia and bulk cell populations were isolated using immunopanning (adapted from ^82,83^).

Following brain collection, meninges and blood clots were manually removed with forceps and brain tissue was processed into small (<1mm^3^) pieces. Tissue was incubated in 10ml papain buffer (20U/ml Papain, Worthington, OH, USA), 0.46% glucose, 26mM NaHCO_3_ 0.5mM EDTA, 1x EBSS) and 200µl DNase I (0.4%, 12,500 U/ml, Worthington) at 34°C for 100 minutes with a consistent CO_2_ flow and periodic shaking.

Digested tissue was firstly washed and then gently triturated in low ovomucoid protease inhibitor solution (0.015 g/ml BSA (Sigma), 0.015g/ml trypsin inhibitor (Worthington) in DPBS at pH of 7.4) to form a single cell suspension. High ovomucoid solution (0.030 g/ml BSA (Sigma), 0.030g/ml trypsin inhibitor (Worthington), in DPBS at pH of 7.4) was gently pipetted to form a layer underneath the cell suspension and cells were spun down for 15 minutes at 250 x g. The cell pellet was resuspended in 0.02% BSA solution and filtered through a nitrex mesh to remove cell clumps and debris. Pre-prepared immunopanning dishes were setup as follows: a 10cm petri dish was incubated at 4°C overnight with 30ul anti-rat antibody (Jackson ImmunoResearch Laboratories, Inc, 112-005-167) in 10 ml 50 mM Tris-HCL (pH 9.5). Following overnight incubation the plate was washed 3 times with DPBS and then incubated with 20 µl anti-CD45 antibody (BD Pharmingen, catalog #550539) in 12 ml of 0.2% BSA for a minimum of 2 hours. Plates were washed 3 times with DPBS and Ssingle cell suspensions were added to pre-coated plates and incubated for 30 minutes with periodic shaking. Unbound cells were collected, spun down and then resuspended in 700µl Qiazol. This population of microglial depleted CNS cells was referred to as ‘Bulk’ and used as a comparison population for downstream analysis. Bound cells (CD45^+ve^ Microglia) were scraped off the plate into 700 µl Qiazol for further processing.

### Library construction and RNA-sequencing

Total RNA was extracted using the miRNeasy mini kit (Qiagen) in accordance with manufacturers protocol. The quality of RNA samples was assessed using a bioanalyzer. Samples with an RNA integrity score (RIN) greater than 7 were used for library construction. One sample did not pass RNA quality control and so was excluded from library preparation. mRNA libraries were prepared using MGIEasy RNA Directional RNA library prep chemistry (MGI), and small RNA libraries were prepared using the MGIEasy Small RNA library prep chemistry (MGI). Libraries were pooled and sequenced across 4 lanes of MGISEQ2000-RS sequencing technology, yielding approximately 400M reads per lane (paired-end 100bp). Library construction and sequencing were performed by personnel at Micromon Genomics (Monash University, Melbourne, Australia).

### Preprocessing and genome alignment of miRNA/mRNA sequencing data

Adapter sequences were removed from small RNA sequencing data and remaining reads were filtered by size (>18 bp) using fastp (ver 0.20.0). Trimmed reads were mapped to the mouse reference genome (mm10/GRCm38) using mirDeep2 with default settings (ver 0.1.3). Candidate novel miRNAs detected by miRDeep2 with a miRDeep2 score > 4 and randfold *p* value < 0.05 were included in downstream analysis. mRNA sequencing data was mapped to the mouse reference genome (mm10/GRCm38) using STAR with default settings (ver 2.7.3). Gene read counts were generated using the featureCounts pipeline in Subread with the following flags (-T 8 -s 2 -p -a) (ver 2.0.0).

### Count normalisation and differential expression analysis

Normalisation and differential expression analysis of miRNA and mRNA was performed using limma (ver 3.48.1, 10.18129/B9.bioc.limma) and edgeR (ver 3.34.0, 10.18129/B9.bioc.edgeR) [53]. Based on multi-dimensional scaling (MDS) analysis one mouse was identified as an outlier and its miRNA/mRNA expression was removed from downstream analysis. miRNAs were filtered for a minimum expression of 1 count per million (CPM) (for miRNA), and 0.5 CPM (for mRNA) in at least 5 samples. Gene counts were normalized using the TMM normalisation method (edgeR). All regression analyses were set up as mixed effects models including age, sex, and cell type (microglia or bulk) as fixed effects and mouse ID was treated as a random effect^84^. The general enrichment analysis was performed by direct comparison of miRNA expression between all microglia and all bulk samples. Specific age enrichment analyses compared only the microglia and bulk samples within a relevant age group. Pairwise analyses of microglial miRNA expression were performed by direct comparison of miRNA expression between microglial age groups. Sex-specific analyses were performed by comparing male and female microglia samples within each age group. For all differential expression analyses, differentially expressed genes were identified using the TREAT method (t-tests relative to a threshold), set at a minimum threshold of a ±2-fold change in expression with a false discovery rate (FDR) < 0.05^85^. Volcano plots were generated using EnhancedVolcano (ver 1.10.0, 10.18129/B9.bioc.EnhancedVolcano)Tables were generated with gt (ver0.3.1) (https://cran.r-project.org/web/packages/gt/index.html).

### miRNA target analysis and miRNA-mRNA regulatory network generation

To infer miRNA-mRNA interactions from gene expression data, Pearson correlations were calculated from normalised counts for all pairs of miRNA-mRNA using psych (ver 2.16, https://CRAN.R-project.org/package=psych). Only those correlations that were strongly negative and significant (r < -0.894, FDR <0.05) were retained for downstream analysis.

To further validate significant negative correlations, miRNA-mRNA interactions were screened for experimental evidence of interaction in any cellular system using multimiR (ver 1.140, 10.18129/B9.bioc.multiMiR) with default settings. For novel candidate miRNAs, mRNA target binding prediction was performed using RNAhybrid (ver 2.2.1, https://bibiserv.cebitec.unibielefeld.de/rnahybrid), whereby novel miRNA sequences were aligned with 3’-UTR mouse sequences (ENSEMBL, release 104) using the following parameters (-e -25 -p 0.05).

miRNA-mRNA interactions which were identified as negatively correlated from expression datasets and validated by multimiR or RNAHybrid were included in downstream pathway analyses and network construction. Visualisations of miRNA-mRNA networks were generated using Cytoscape (https://cytoscape.org/). KEGG (Kyoto Encyclopedia of Genes and Genomes) and GO (gene ontology) enrichment analysis were performed on the target mRNAs in generated networks using the kegga and goanna modules in limma respectively (ver 3.48.1, 10.18129/B9.bioc.limma).

### qRT-PCR of mRNA expression

100ng of total RNA isolated from P6 microglia/bulk samples was reverse transcribed using Taqman reverse transcription reagents (Invitrogen) in accordance with manufacturer’s instructions. 25 ul qPCR reactions were prepared as follows; 12.5 µl 2x SYBR green master mix (ABI Biosystems, MA, USA), 1µM of forward and reverse primers and DNAse free water. Primer sequences are outlined in Table 5. Expression of 18S was used to normalise for RNA input. Relative fold change of expression was calculated using the 2(-delta-delta C(T)) method and log transformed^86^.

**Table 5:**
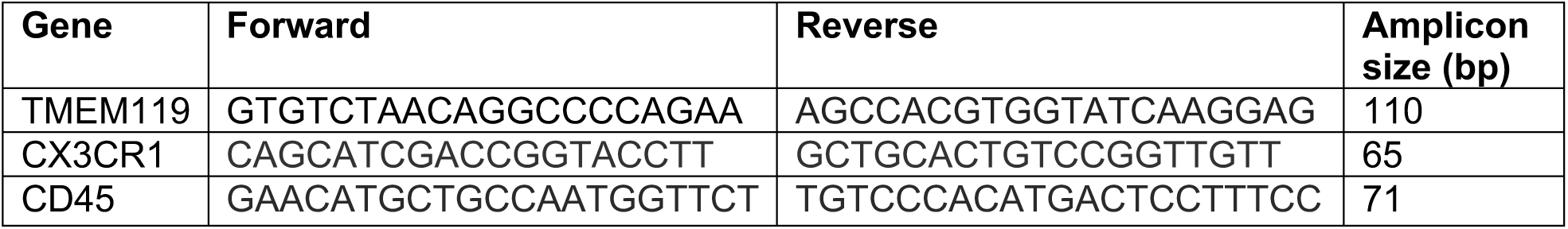
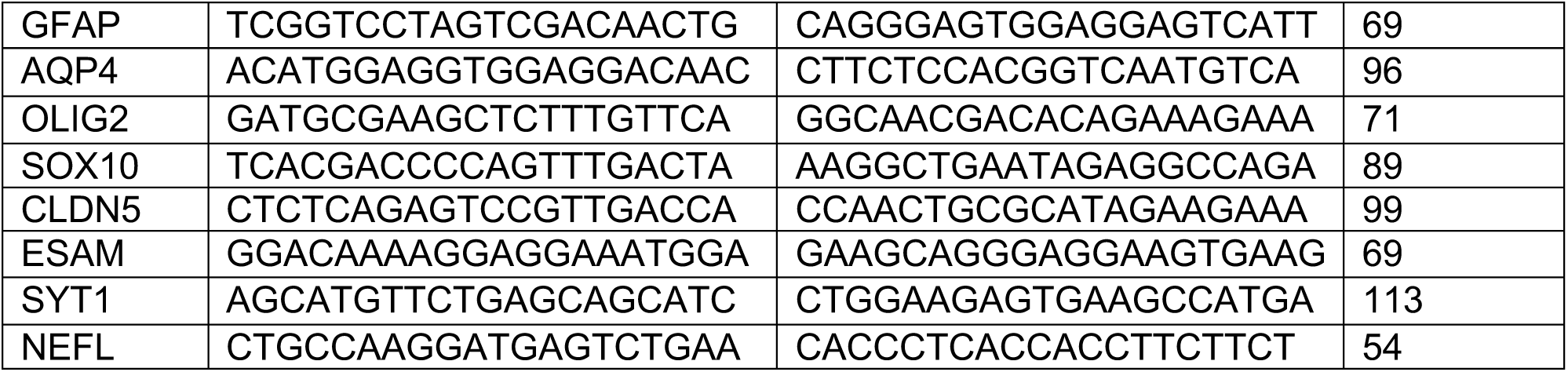
Primer sequences for mRNA qPCR of immunopanning samples

### qRT-PCR of miRNA expression

50ng of total RNA isolated from microglia/bulk samples was reverse transcribed using the miRCURY LNA RT kit (Qiagen) in accordance with manufacturer’s instructions. The miRCURY LNA miRNA PCR assay system (Qiagen) was used to detect specific miRNA expression. Expression of ribosomal 5S was used to normalise for RNA input for all samples[56]. Relative fold change of expression was calculated using the 2(-delta-delta C(T)) method and log transformed. Paired t-tests were performed to identify consistent expression differences between paired microglia/bulk samples for each miRNA. Unpaired t-tests were performed to identify differences in candidate miRNA expression between male and female microglia subgroups. Statistical analyses were performed using GraphPad Prism (ver 9.2.0, Graphpad, CA, USA).

## Supplementary Figures

**Figure S1:**
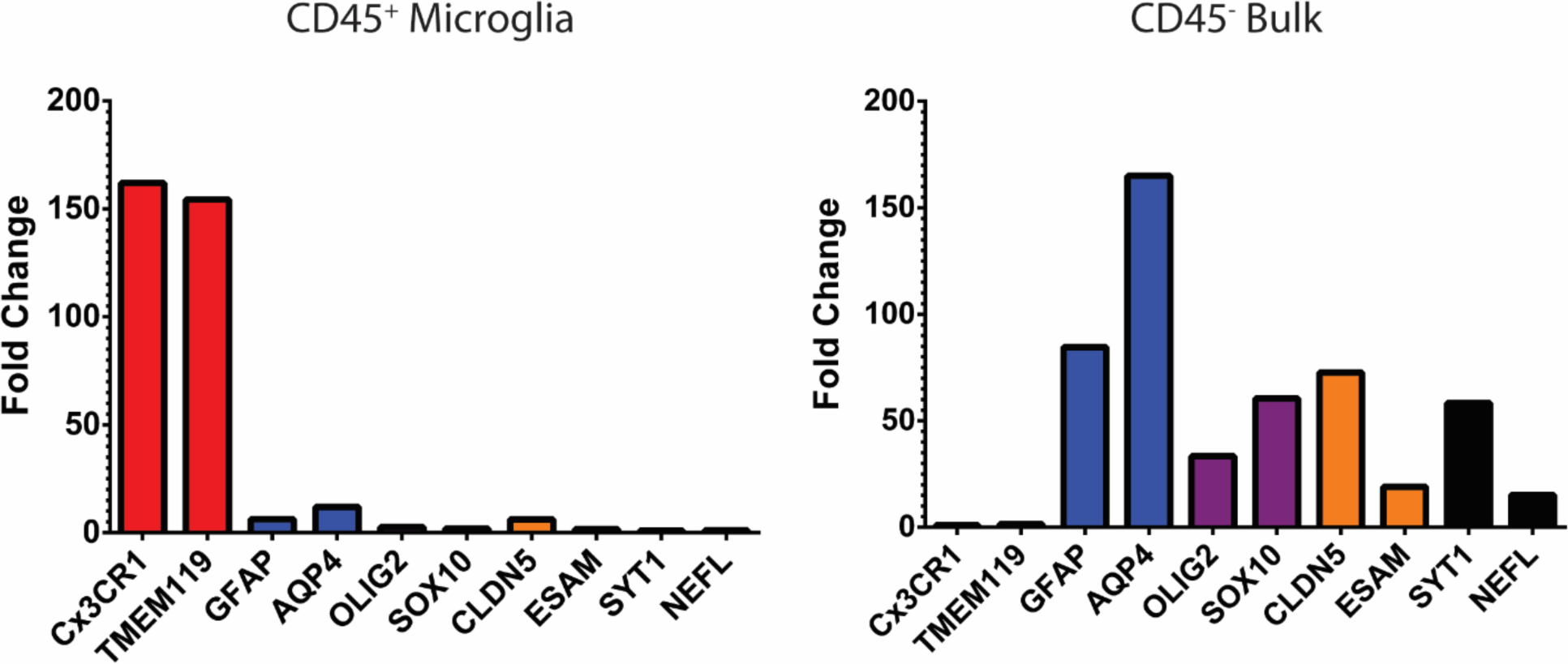
Validation of CD45^+ve^ immunopanning technique via qPCR. qPCR of 10 target genes from CD45^+ve^ and CD45^-^ immunopanning samples from a P6 mouse. Gene targets represent major cell types of the CNS; Microglia (CX3CR1,TMEM119), Astrocytes (GFAP, AQP4), Oligodendrocytes/OPCs (OLIG2, SOX10), Endothelial Cells (CLDN5, ESAM), Neurons (SYT1, NEFL). CD45^+^ populations exhibit strong enrichment of microglial gene targets and low contamination of other cell types. Conversely, the CD45^-ve^ sample exhibits expression of all other cell types and very low contamination of microglial cells.

## Data availability statement

The data that support the findings of this study are openly available in the gene expression omnibus (GEO) at [https://www.ncbi.nlm.nih.gov/geo/], reference number [*reference number will be provided upon acceptance of the manuscript*].

## Acknowledgements

The authors would like to acknowledge the support of the Australian Government Research Training Program Scholarship (ADW), National Health and Medical Research Council (APP1175775; TJK), Multiple Sclerosis Australia (21-6-008; TJK and 21-3-038; SS). The Florey Institute of Neuroscience and Mental Health and the Walter and Eliza Hall Institute of Medical Research acknowledge the strong support from the Victorian Government, particularly, the funding from the Operational Infrastructure Support Grant.

## Author contributions

Conceived and designed the experiments: MDB TJK AW BAA. Performed the experiments: ADW SS AA MDB. Analysed and interpreted the data: ADW SAS TJK BAA MDB. Wrote the paper: ADW MDB BAA.

## Competing interests (mandatory)

The authors declare no competing interests.

